# Multimodal analysis of osteoarthritic chondrocytes reveals mitochondrial alterations and patient-specific OxPhos response to bezafibrate

**DOI:** 10.1101/2025.10.28.684789

**Authors:** Lucie Danet, Romain Guiho, Sally Hussein, Anaïs Defois, Nicolas Gaigeard, Joëlle Veziers, Guillaume Mabilleau, Antoine Hamel, Denis Waast, Jérôme Guicheux, Marie-Astrid Boutet, Claire Vinatier

**Affiliations:** Nantes Université, Oniris, INSERM, CHU Nantes, UMR1229 Regenerative Medicine and Skeleton, RMeS, Nantes, France; Nantes Université, Univ Angers, Oniris, CHU Nantes, INSERM, Regenerative Medicine and Skeleton, RMeS, UMR 1229; CHU Angers, Departement pathologie cellulaire et tissulaire, Angers, France; Centre for Experimental Medicine & Rheumatology, William Harvey Research Institute and Barts and The London School of Medicine and Dentistry, Queen Mary University of London, EC1M 6BQ, London, United Kingdom

**Author notes:** Corresponding author: Claire Vinatier.

**Keywords:** Osteoarthritis, Cartilage, Chondrocytes, Mitochondria

## Abstract

**Background:** Osteoarthritis (OA) is the most common joint disease and is characterized by bone remodeling, cartilage degradation and synovial inflammation. To date, no effective treatment is available for this debilitating condition. Recent evidence suggests that mitochondrial dysfunction, including oxidative phosphorylation (OxPhos) failure, accumulates within OA chondrocytes and may contribute to pathogenesis. In this context, mitochondrial dysfunction may be associated with observable changes in mitochondrial number, size and shape. However, a comprehensive characterization of mitochondria-related features during OA, from tissue-to-cell level, is still lacking. Addressing these gaps could inform therapeutic strategies, such as the partial restoration of OxPhos, which has been proposed as a therapeutic approach.

**Methods:** Here, we employed a multimodal approach that included Fourier-transform infrared spectroscopy (FTIR), scanning transmission electron microscopy (STEM) and real-time cellular metabolic assays (Seahorse technology) to better characterize mitochondrial parameters in cartilage during OA. Two types of experimental models were used using human cartilage: (1) undamaged versus damaged OA zones, and (2) non-OA versus OA samples. In addition, we investigated the potential of repurposing bezafibrate, an approved peroxisome proliferator-activated receptor (PPAR) agonist, as a mitochondria-based therapy to restore OxPhos in OA chondrocytes.

**Results:** We identified that OA chondrocytes exhibit a decrease in glycogen deposits surface, and an increased number of mitochondria alongside an OxPhos dysfunction compared to non-OA chondrocytes. A similar trend toward glycogen storage deficiency and increased mitochondria number was observed in OA chondrocytes from damaged cartilage areas. Furthermore, multivariate analyses revealed that the clinical profiles of OA patients allowed OA chondrocytes to be separated into responders and non-responders to bezafibrate.

**Conclusion:** We provide evidence that OA chondrocytes display decreased glycogen deposits surface, increased mitochondrial number and OxPhos dysfunction. Additionally, we identified that bezafibrate, a PPAR agonist, improved OxPhos function in a subgroup of OA chondrocytes derived from patients.

## INTRODUCTION

Osteoarthritis (OA) is the most prevalent joint disease, affecting over 500 million people worldwide, predominantly among the elderly population [1]. This condition is mainly characterized by progressive cartilage degradation, subchondral bone remodeling and osteophyte formation, as well as synovitis. However, no curative treatment is currently available. The management of the disease remains limited to symptomatic relief, ranging from non-pharmacological approaches (e.g., weight loss and physical exercise) to pharmacological interventions (e.g., non-steroidal anti-inflammatory drugs and analgesics), and surgical procedures (total joint replacement) for end-stage OA [2]. Therefore, OA management places a heavy socio-economic burden on the healthcare systems, making the search for disease-modifying osteoarthritis drugs (DMOADs) crucial [3,4].

OA has long been regarded as a “wear and tear” disorder. However, cumulative evidence indicates that OA is far more complex, characterized by high heterogeneity and multifactorial origins. Various classification attempts have been proposed, such as clinical phenotypes and molecular endotypes [5–7]. In parallel, aging, which remains the main risk factor for developing OA and thus a central contributor of the disease pathophysiology, is linked to several established biological hallmarks notably mitochondrial dysfunction [8].

Recent evidence describes that OA chondrocytes exhibit mitochondrial dysfunction, which can contribute to disease pathogenesis [9–11]. These alterations occur at multiple levels. Mitochondrial dynamics are disrupted in human OA chondrocytes, resulting in a fragmented mitochondrial network driven by active fission processes over fusion. This phenomenon is further exacerbated in pro-inflammatory environments [12]. Mitochondrial biogenesis, the process of synthesizing new mitochondria, is also impaired, as evidenced by the downregulation of PGC-1α, the master regulator of this pathway [13]. Additionally, mitophagy, the selective removal of damaged mitochondria, which is regulated in part by Parkin and PINK1, is altered in OA chondrocytes [14–16]. Dysfunctional mitochondria therefore accumulate and exert harmful effects on chondrocytes. All of the above mitochondrial features are structural and can therefore influence mitochondrial abundance. Nevertheless, there is still no consensus on whether the number, size, and shape of mitochondria are altered in OA chondrocytes.

One main function of the mitochondria is energy metabolism. Although chondrocytes reside in a hypoxic environment and generate most of their ATP through glycolysis (∼75%), oxidative phosphorylation (OxPhos) remains active and also generates ATP (∼25%), thereby supporting cartilage anabolism [17]. In OA chondrocytes, OxPhos dysfunction has been reported, including reduced activity and expression of mitochondrial respiratory chain complexes [13,18]. While OA cartilage undergoes neovascularization and can be more easily supplied with oxygen, a counterintuitive phenomenon similar to the Warburg effect can occur in chondrocytes, promoting aerobic glycolysis over OxPhos [19,20]. Additionally, a high level of glucose can inhibit OxPhos through the Crabtree effect. OA chondrocytes shift their metabolism to rely more on glycolysis, as illustrated by the high expression of glycolytic enzymes such as pyruvate kinase M2 (PKM2) [21] and lactate dehydrogenase A (LDHA) [22], and by the increased levels of glycolysis-related metabolites in synovial fluid. In this context, we and others have demonstrated that OA chondrocytes undergo a shift toward using less OxPhos for further glycolysis, which can promote pro-inflammatory and catabolic processes [23–25].

Given this mitochondrial dysfunction, proof-of-concept studies have investigated the effect of intra-articular transplantation of mitochondria in OA animal models and showed that it can ameliorate disease progression [26–28]. More specifically, the sole intra-articular delivery of high-energy photosynthetic units was sufficient to reproduce this phenotype, highlighting mitochondrial energetic dysfunction as a promising therapeutic target for OA [29]. However, their long-term efficacy and safety must be demonstrated before considering clinical translation. Therapies addressing mitochondrial functions, and more specifically, OxPhos stimulation therapies for OA chondrocytes, could logically be considered as first-line approach. Nevertheless, a comprehensive characterization of glucose metabolism and mitochondrial parameters in OA cartilage compared to non-OA chondrocytes, both at the tissular and cellular levels, is still lacking. Furthermore, it is unclear whether pathological changes in mitochondrial abundance, size, and shape occur and if these changes are associated with OxPhos dysfunction in OA chondrocytes.

The primary goal of this study was to better characterize mitochondrial features in OA chondrocytes (OAC) using a multimodal approach enabling a multiscale analysis from tissue to the cellular level. We employed Fourier-transform infrared spectroscopy (FTIR), scanning transmission electron microscopy (STEM), and real-time cellular metabolic assays in two distinct ex vivo and in vitro human OA chondrocyte experimental models. This study also aimed to investigate the potential of repurposing the approved peroxisome proliferator-activated receptor (PPAR) agonist, bezafibrate, to improve OxPhos function in OAC.

## MATERIAL AND METHODS

### OA patients

Clinical data were collected for each OA patient included in this study. Sex, age, BMI and Kellgren-Lawrence score of the OA patients grouped according to the technique in which they were included are provided in **Supplementary Table 1**.

### Cartilage samples

Cartilage pieces from 9 non-OA patients and 31 OA patients were obtained as surgical waste, respectively, from the vertebral transverse costal facet of scoliotic patients, and from the femoral condyles and tibial plateaus of knee joints of patients undergoing total knee arthroplasty for end-stage OA. Written informed consent was obtained from all patients in accordance with the Declaration of Helsinki. The study protocol was approved by the committee for the protection of Persons Pays de la Loire and the French Ministry of Higher Education and Research (registration number: DC-2017–2987).

### Histological analyses

Tibial plateaus from OA patients were fixed in 4% paraformaldehyde (PFA) for one week and then cut into small pieces. Samples were decalcified in 0.636M ethylenediaminetetraacetic acid (EDTA, 0.636 mol/L, pH 7.5–8) at 4°C, dehydrated and embedded in paraffin. Paraffin-embedded OA cartilage samples from both undamaged and damaged zones of the tibial plateaus were sectioned at 4 µm thickness for further analysis.

#### OARSI scores

The characterization of cartilage degradation was performed using the Osteoarthritis Research Society International (OARSI) score [30] on 4-µm-thick sections stained with Safranin O/Fast Green. All sections were digitally scanned using Nanozoomer S360 (Hamamatsu Photonics).

#### FTIR microscopy

Sections of OA cartilage were deposited on barium fluoride (BaF2) windows (Crystran Ltd). Spectra were acquired using an Invenio-S spectrometer coupled to a Bruker Hyperion II infrared microscope equipped with a 64 × 64 focal plane array (FPA) detector. This infrared microspectrometer was calibrated daily to ensure accuracy in wavenumber and absorbed energy. Several regions of interest (ROI) of at least 1.2 mm width spanning the full articular cartilage depth were analyzed with a 15X Cassegrain objective (NA = 0.4). Spectra were collected in transmission mode in the mid-infrared range (850–2000 cm^-1^) with 32 accumulations at 4 cm⁻¹ resolution. Paraffin/BaF2 windows background spectra were also recorded within the same conditions. Data processing using Matlab (version R2025a, MathWorks) included subtraction of paraffin signal at ∼1465 cm⁻¹, normalization to the Amide I band (∼1655 cm^−1^), and a denoising using the Savitzky-Golay algorithm (degree 2, window size 9). Quality control of each spectrum was conducted by calculating the signal-to-noise ratio (SNR) within the spectral range of 1850– 2000 cm⁻¹, which is free of biological signals. Spectra with an SNR ≥10 were considered sufficient to proceed with further analyses. Second derivatives were computed over the spectral range 1034–1185 cm^−1^ and used as a loading vector for curve fitting. Spectral curve fitting quality was assessed by root mean square error, with a threshold set at 1% of the total spectral interval area. Intensities of sub-bands located at ∼1062 cm^-1^, ∼1152 cm^−1^, and ∼1127 cm^−1^ normalized to the ∼1655 cm^−1^ amide I band were used to estimate the proteoglycan (PG), glycogen (GLY), and lactic acid (LA) contents, respectively [31–33].

### Scanning transmission electron microscopy (STEM)

3 non-OA cartilage and 5 OA samples (including 3 with both undamaged (U) and damaged (D) regions from the same OA joints) were cut into small pieces approximately 1 mm thick and 0.5 cm wide for scanning transmission electron microscopy (STEM) preparation. Samples were fixed overnight at 4°C in 1.6% glutaraldehyde (R1011, agar scientific), in 0.1 M phosphate buffer pH 7.5 (21180-21190, EMSdiasum), rinsed, and post-fixed for 1 hour in 1% osmium tetroxide (R1021, agar scientific), and 1.5% potassium ferrocyanide (P9387, Sigma-Aldrich) in ultrapure water, at room temperature and protected from light. Then, the samples were rinsed with ultrapure water and dehydrated through a graded series of ethanol and acetone solutions, infiltrated with increasing concentrations of Epon in acetone, using the agar 100 resin kit (R1031, agar scientific), and embedded in flat molds. After polymerization in a dry oven at 37°C and then 60°C, the resin-embedded blocks were trimmed and cut into 700 nm thick sections to detect zones of interest containing cells using toluidine blue and azur blue staining. Then, the sections were cut into 80-nm-thick ultrathin sections for STEM observations. Imaging was performed using a STEM detector on a Cryo Scanning Electron Microscope (GeminiSEM 300, ZEISS), using an accelerating voltage of 25-30 kV with a 7.5 µm diameter aperture. Image pixel size was 3.7 nm. Image analysis was conducted in QuPath (version 0.5.1). Mitochondria were manually delineated by 4 independent readers, and quantitative parameters, including number, size, area, solidity, Feret diameters, and aspect ratio were extracted. Solidity is defined as the ratio of the mitochondrial area to the area of its convex hull. Aspect ratio, defined as the ratio of the major axis (maximum Feret diameter) to the minor axis (minimum Feret diameter) of an object, was used to assess mitochondrial shape elongation. Information on glycogen deposits was also extracted. OA chondrocytes from D regions were compared with those from U regions, and OA chondrocytes (OAC) were compared to non-OA chondrocytes (NC).

### mtDNA copy number quantification

Genomic DNA (gDNA) was extracted from both non-OA and OA cartilage pieces using the PureGene Core kit A (1042601, Qiagen). Cartilage explants were dissociated in 300 µL of cell lysis solution, treated with 15 µL of proteinase K (20 mg/mL, EU0090-B, Euromedex), and incubated overnight at 56°C. gDNA was purified according to the manufacturer’s protocol, and quantified using a Nanodrop spectrophotometer. Real-Time quantitative PCR was prepared using SYBR Select Master Mix (4472908, Thermo Fisher Scientific) and then performed using a CFX96™ Touch Deep Well Real-Time PCR Detection System (1855195, Bio-Rad). Primer sequences are listed in **Table 1**. The ND1 gene serves as a mitochondrial target, while GAPDH was used as a nuclear reference gene.

**Table 1.**
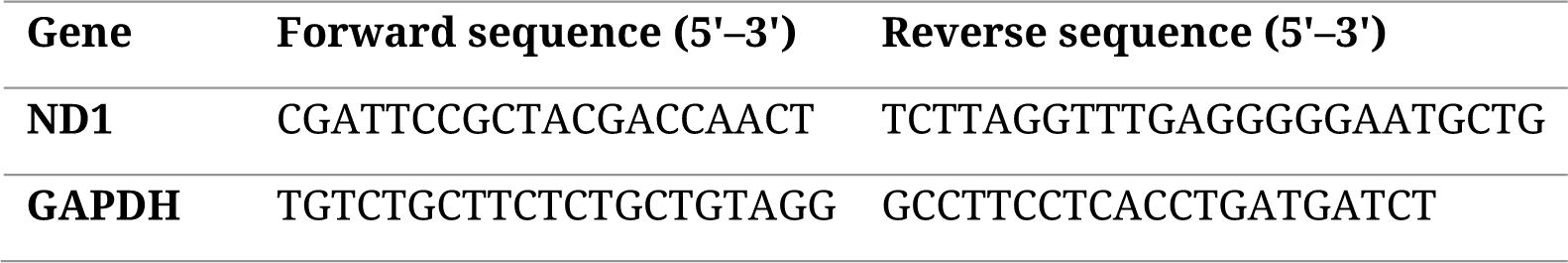
Primers used for mtDNA copy number quantification.

### Primary human chondrocytes isolation and culture

Primary human chondrocytes were isolated from surgical waste cartilage as described in the cartilage samples section, primarily from the femoral condyles. Non-OA cartilage and OA cartilage were cut and digested as previously described [25]. Briefly, small cartilage pieces were digested in two steps: first in 0.2% collagenase (125 U/mg, C6885, Sigma-Aldrich) for 30 min at 37°C in Hanks’ Balanced Salt Solution (HBSS, L0606-500, Biowest) supplemented with 10% penicillin-streptomycin solution (PS, 1000 U/mL, 15140122, Thermo Fisher Scientific) and then in 0.03% collagenase in complete medium (high-glucose Dulbecco’s Modified Eagle’s Medium, DMEM, 31966021, Gibco) with 10% fetal bovine serum (FBS, Dominique Dutscher) and 1% PS for approximately 14 h at 37°C. Digestion products were filtered through a 70 μm strainer and centrifuged at 280 g for 3 min to remove debris. NC and OAC cultured cells were seeded at a density of 1.5 × 10⁴ cells/cm², and 6.0 × 10⁴ cells/cm², respectively, and maintained in complete medium at 37°C in a humidified atmosphere with 5% CO₂, with medium changes three times per week. Chondrocytes were used exclusively at passage 1.

### Real-time cellular metabolic assays in Seahorse

NC and OAC chondrocytes were seeded at 60,000 cells/well in Seahorse XF96 microplates (103794-100, Agilent) for both ATP Rate Assay and Mito Stress Test. After 24 h, OAC were treated with 0.1, 1 or 10 µM bezafibrate (B727, Sigma-Aldrich) or vehicle (DMSO) as a control for an additional 24 h. Before analysis, culture medium was replaced with phenol red-free DMEM (D5030, Sigma-Aldrich) supplemented with 4 mM glutamine, 25 mM glucose, and 1 mM pyruvate (pH 7.4). Microplates were incubated for 45 min at 37°C in a CO2-free environment. The sensor cartridge was hydrated in Seahorse XF calibrant (100840-000, Agilent) at 37°C for at least 24 h in a humidified environment, and then loaded as follows: 25 µL per port of oligomycin (2 µM), carbonylcyanide-3-chlorophenylhydrazon (CCCP, 4 µM), and a 1:1 mixture of rotenone and antimycine A (1 µM each) for the Mito Stress Test. For the ATP Rate Assay, 25 µL per port of oligomycin (2 µM) and the 1:1 rotenone/antimycin A mixture (1 µM each) were loaded. Oxygen consumption rate (OCR) was measured every 5 min before and after the successive drug injections using the Seahorse XF Pro analyzer (Agilent). Data were analyzed using Wave software (Agilent).

### Partial Least Squares-Discriminant Analysis (PLS-DA)

Based on Seahorse Mito Stress Test results, patient-derived OAC were retrospectively categorized as responders or non-responders to bezafibrate treatment. PLS-DA was performed using the scikit-learn package in Python (version 3.13.6), to identify clinical variables discriminating between responders and non-responders groups.

### Statistical analyses

All statistical analyses, except for PLS-DA, were performed using GraphPad Prism software (version 8.0). Paired data were analyzed using the Wilcoxon signed-rank test, and unpaired data were analyzed using the Mann-Whitney test, as indicated in the figure legends. Correlation analyses were conducted using the Spearman correlation test. For comparisons of clinical data, the Kruskal–Wallis test was applied for numerical variables, and Fisher’s or Chi² test were used for categorical variables, depending on sample size. A p-value < 0.05 was considered statistically significant (*p < 0.05). All graphs, except the radar plot, were generated using GraphPad Prism software (version 8.0). The radar plot graph was generated using R (version 4.5.1) on RStudio (version 2025.05.1+51).

## RESULTS

### Investigation of glucose metabolism and mitochondrial features in damaged compared to undamaged OA cartilage

Given that energy metabolism is known to be dysregulated in OA chondrocytes, we first assessed whether components of glucose metabolism were altered at the tissue level before investigating mitochondrial features. 6 matched samples of visually undamaged (U) and damaged (D) cartilage zones from OA patients were used. The overall experimental workflow is illustrated in **Figure 1A**. Undamaged cartilage displayed a thicker layer of chondrocytes and smooth articular surface, whereas damaged cartilage showed a thinner structure, reduced Safranin O staining, and surface irregularities (**Supplementary Figure 1**). OARSI scores were significantly higher in damaged zones (4.97 ± 0.20) compared with undamaged zones (2.80 ± 0.93), confirming cartilage degradation and supporting the validity of this model to assess OA progression through structural damage (**Figure 1B**). We applied a supervised analysis targeting specific molecular entities relevant to cartilage and glucose metabolism, selected for their well-defined absorption bands reported in the literature: proteoglycan, glycogen, and lactic acid. Representative images are presented in **Figure 1C** and in **Supplementary Figure S2**. The two sets of images are displayed at different scales. In undamaged OA, the cartilage was thicker, requiring a larger ROI, whereas in damaged OA, only a thin layer remained, leading to the use of a smaller ROI. Quantitative analyses revealed a high interpatient variability regarding their changes in proteoglycan and glycogen content in damaged compared to undamaged OA cartilage zones, with no significant changes observed across conditions (**Figure 1D**). However, when all cartilage zones were considered independently of their previous damaged or undamaged classification, FTIR analysis revealed a significant negative correlation between proteoglycan content and OARSI score (**Figure 1E**). Interestingly, glycogen content also showed an inverse correlation with OARSI scores, suggesting a progressive loss of intracellular glycogen in chondrocytes with increasing disease severity (**Figure 1E**). In contrast, lactic acid levels remained unchanged with respect to OARSI scores (**Supplementary Figure 2**).

**Figure 1.**
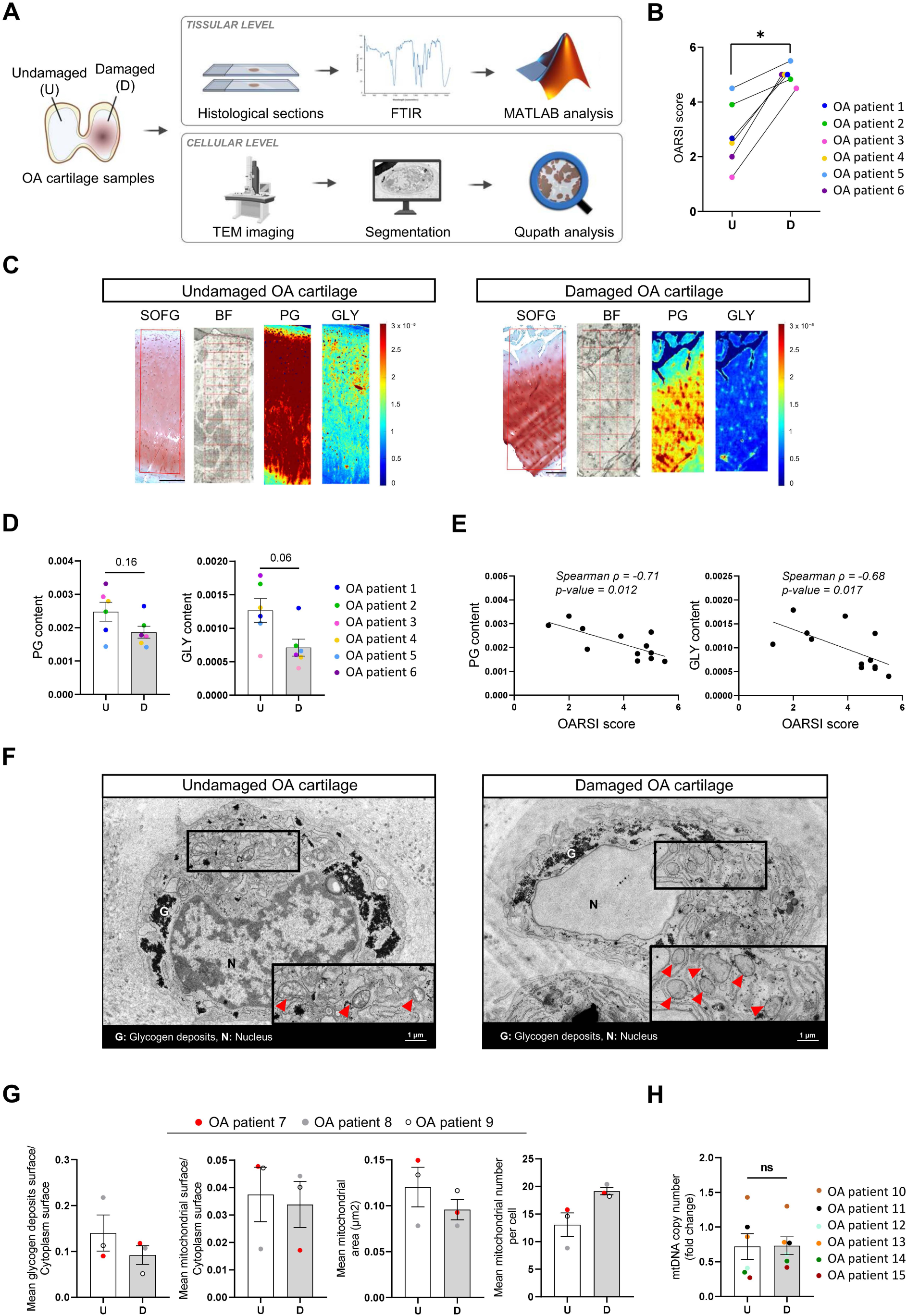
Investigation of glucose metabolism and mitochondrial features in damaged compared to undamaged OA cartilage using FTIR and STEM. **A.** Overview of the experimental workflow. Undamaged (U) and damaged (D) OA cartilage samples were obtained from distinct areas within the same OA human knee joint. **B.** OARSI scores of paired OA cartilage samples (N=6). **C.** FTIR spectroscopic imaging of proteoglycan (PG) and glycogen content (GLY) (N=6). Safranin O/Fast Green (SOFG) staining was used to guide the selection of regions of interest (ROI) in bright field (BF) images for the FTIR analysis. The images shown correspond to the OA patient 4. Scale bar, 250 µm for undamaged and 500 µm for damaged OA cartilage. **D.** Quantification of proteoglycan and glycogen content normalized to the amide I band measured by FTIR spectroscopy (N=6). Data are represented as the mean experimental spatial values from each patient’s D or U zones. **E.** Spearman correlation between proteoglycan and glycogen content measured by FTIR in cartilage zones and OARSI scores. **F.** Representative STEM images of OA chondrocytes within cartilage explants from U and D zones. Glycogen deposits (G), nuclei (N), and mitochondria (red arrows) are indicated. Enlarged views highlight mitochondrial morphology. **G.** Quantification of glycogen and mitochondrial parameters from STEM images (N=3, n ≥ 6 cells). **H**. mtDNA copy number presented as fold-change using the undamaged group as a reference (N=6). *p < 0.05, ns, non-significant, p-values were calculated using the paired Wilcoxon signed-rank test comparing undamaged and damaged OA cartilage groups [B, D, and H]. Each dot represents a patient. Values are expressed as means ± SEM [D, G, and H]. The Spearman correlation test was used, and the corresponding p-value and Spearman’s correlation coefficient (ρ, rho) are indicated on the graph E.

Moving beyond tissue-level metabolic analyses, we then examined the ultrastructure of OA chondrocytes within cartilage explants to visualize glycogen deposits and mitochondria, to assess potential differences between chondrocytes from undamaged and damaged OA cartilage. To this end, 3 paired human OA cartilage samples were analyzed by STEM to image chondrocytes in situ. Representative images of chondrocytes from undamaged and damaged zones are shown in **Figure 1F**. Subcellular granular black aggregates typical of glycogen deposits were identified and mitochondria were recognized by their spherical or elongated shape, the presence of cristae, and their double membrane (indicated by red arrows). Quantitative analyses revealed that the proportion of cytoplasmic surface occupied by glycogen showed a tendendy to be lower in chondrocytes from damaged zones compared with undamaged zones (**Figure 1G**). While the mean mitochondrial surface area did not differ markedly between zones, this average concealed strong inter-patient variability, sometimes with opposite trends (**Figure 1G**). Finally, across all patients, mitochondrial size appeared reduced and mitochondria number increased in chondrocytes from damaged zones, although not reaching significant differences in mitochondrial DNA (mtDNA) copy number (**Figure 1H**). From STEM images, quantification of glycogen deposits and mitochondrial parameters at the per-cell level are represented for individual patients in **Supplementary Figure 3**. Additionally, no changes in the morphological parameters of mitochondrial solidity and aspect ratio were observed between the two groups (**Supplementary Figure 4**). Collectively, these findings suggest a trend toward glycogen deposits decline and a higher number of mitochondria in chondrocytes from damaged compared to undamaged OA cartilage.

### Quantification of glycogen deposits, presence of mitochondria, and OxPhos function in OAC compared to NC

To further characterize mitochondrial alterations linked to OA pathological processes, we expanded our analysis to include non-OA cartilage and compared it with OA cartilage samples. Mitochondrial features were assessed either as structural parameters in non-isolated chondrocytes using STEM imaging, or as functional parameters by assessing energy metabolism in isolated and cultured chondrocytes using real-time cellular metabolic assays. The overall experimental workflow is represented in **Figure 2A**. Representative STEM images of non-OA chondrocytes (NC) and OA chondrocytes (OAC) in their respective explants are shown in **Figure 2B**. Consistent with what observed in damaged versus undamaged OA cartilage, OAC displayed a significant reduction in glycogen deposits compared with NC, suggesting an altered glucose metabolism (**Figure 2C**). In addition, the mitochondrial surface area was increased in OAC, and was not related to changes in mitochondrial size but rather to the increased number of mitochondria. This change was accompanied by a significant increase in mtDNA copy number compared with NC (**Figure 2D**). Again, a high inter-patient variability was observed (**Supplementary Figure 5**). Of note, mitochondrial morphology parameters were similar between groups (**Supplementary Figure 4**).

**Figure 2.**
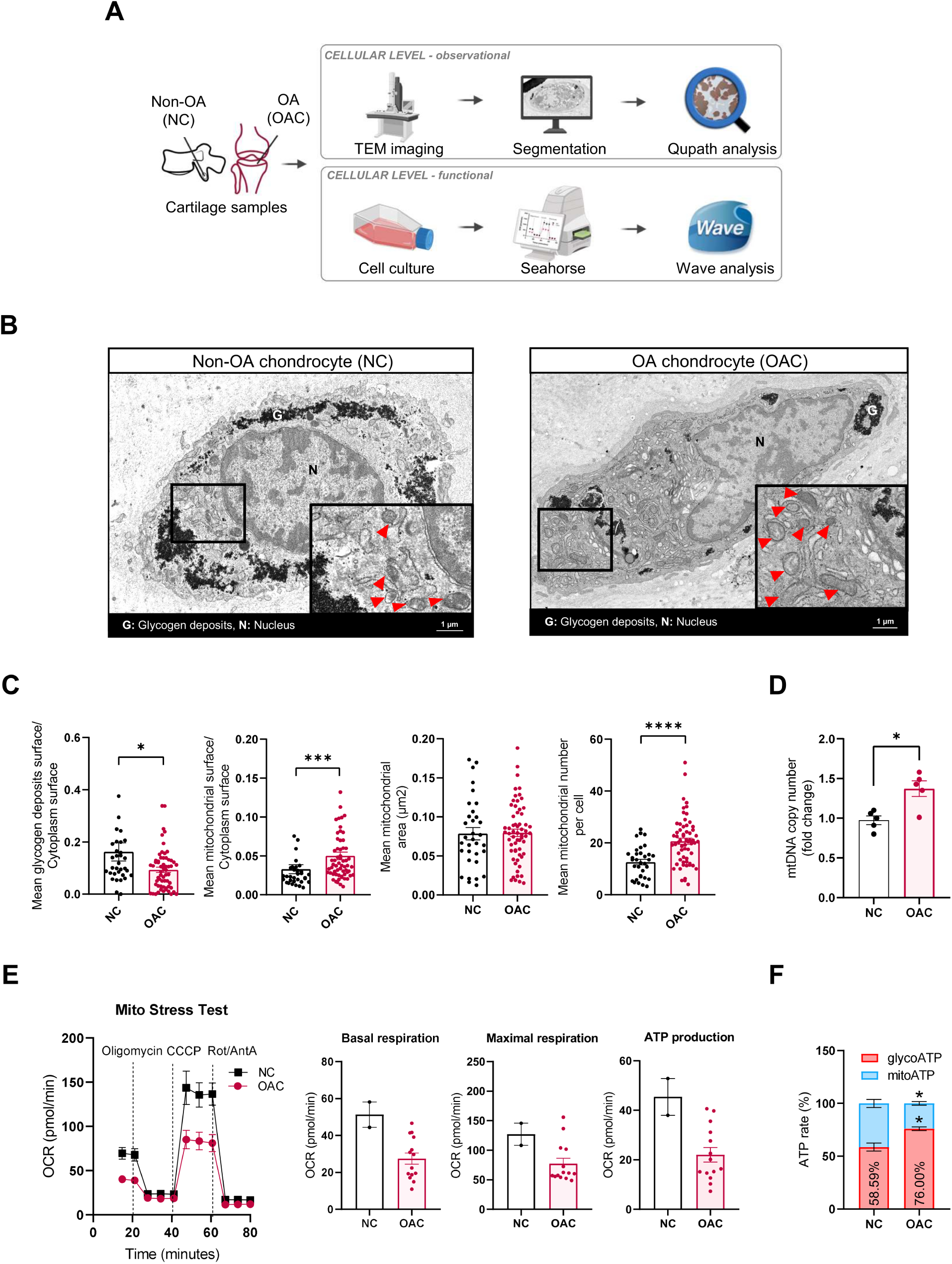
Quantification of glycogen deposits, presence of mitochondria, and OxPhos function in OAC compared to NC using STEM and real-time cellular metabolic assays. **A.** Overview of the experimental workflow. NC and OAC were analyzed either as native cells within cartilage explants [B-D] or as isolated and cultured cells [E-F]. **B.** Representative STEM images of NC and OAC within cartilage explants. **C.** Quantification of glycogen deposits and mitochondrial parameters from STEM images (NC: N=3 patients; OAC: N=5 patients; n≥ 6 cells per patient). Each dot represents a cell. **D.** mtDNA copy number presented as fold-change using NC group as a reference (NC: N=3, OAC: N=5). **E.** Functional analysis of mitochondrial respiration in NC and OAC. OxPhos function was assessed using a Mito Stress Test (Seahorse). Oxygen consumption rate (OCR) values were used to quantify basal respiration, maximal respiration, and ATP production (NC: N=2; OAC: N=14). Each dot represents a patient [D-E]. **F.** Relative contribution of glycolysis (glyco) and OxPhos from the mitochondria (mito) to ATP production assessed by the ATP Rate Assay (Seahorse) (NC: N = 4; OAC: N = 5). *p < 0.05, **p < 0.01, ***p < 0.001, ****p < 0.0001, Mann-Whitney test comparing NC and OAC for panels C, D and F. Values are expressed as means ± SEM.

We next investigated whether the reduction in glycogen level and the increase in mitochondrial number in OAC were associated with functional changes in energy metabolism, focusing on OxPhos activity, a key process responsible for high-yield ATP production by the mitochondria. OAC displayed a trend toward impaired OxPhos compared with NC. Indeed, the real-time cellular metabolic assays that specifically assess the OxPhos function (Mito Stress Test) revealed lower basal respiration, maximal respiration, and ATP production in OAC (**Figure 2E**). To confirm that OxPhos dysfunction affected the source of ATP molecules produced in OAC, we performed a dedicated real-time cellular metabolic assay called ATP Rate Assay. These results demonstrated a significant metabolic shift in OAC, with reduced ATP production from OxPhos (mitoATP) and increased ATP from glycolysis (glycoATP) compared with NC (**Figure 2F**). Thus, OAC preferentially shifted from OxPhos to glycolysis for ATP generation.

### Effect of bezafibrate on OAC OxPhos function

Given that OAC were characterized by impaired OxPhos, with a reduced contribution to overall ATP production, we then tested the effect of bezafibrate, a pan-peroxisome proliferator-activated receptor (PPAR) agonist, on OAC OxPhos function. Bezafibrate has indeed previously been shown to increase ATP production in OxPhos-deficient fibroblasts and reduce ROS levels in neural cells [34,35].

OAC were treated for 24 h with increasing doses of bezafibrate and OxPhos function was then assessed using a Mito Stress Test. When pooling all patient-derived OAC, the average OCR profiles were similar between untreated and bezafibrate-treated groups, with no detectable changes in basal respiration, maximal respiration, or ATP production (**Supplementary Figure 6**). Indeed, bezafibrate did not influence these parameters in half of the OAC that were therefore classified as non-responders (**Figure 3A**). Interestingly, the other half of the samples presented a significant increase in maximal respiration and ATP production upon bezafibrate treatment, particularly at 1µM, and were classified as responders (**Figure 3B**).

**Figure 3.**
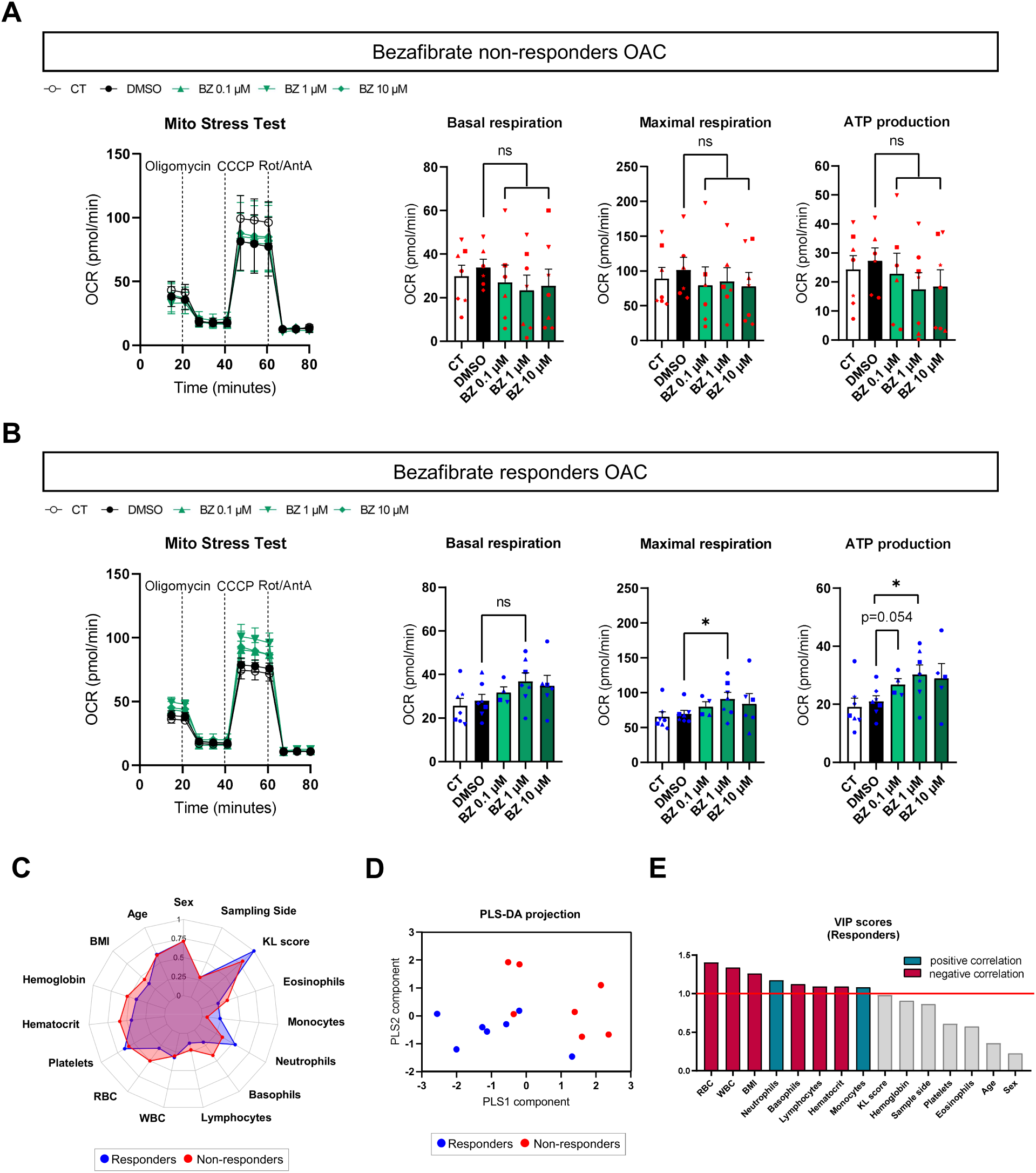
Characterization of OAC response to bezafibrate by integrating real-time cellular metabolic assay and clinical data. (A-B). Functional analysis of mitochondrial respiration in OAC after 24 h treatment with bezafibrate (0.1, 1 or 10 µM). OxPhos function of bezafibrate non-responders (**A**) and responders (**B**) OAC assessed using a Mito Stress Test (Seahorse). Oxygen consumption rate (OCR) values were used to quantify basal respiration, maximal respiration, and ATP production (Total: N=14, Responders: N=7, Non-responders: N=7). (**C-E**). Integration of clinical data from OA patients with responders OAC (blue, N=7) and non-responders OAC (red, N=7). **C.** Radar plot. **D.** PLS-DA projection. **E.** VIP scores for responders OAC derived from PLS-DA analysis. *p < 0.05, ns, non-significant. p-values were calculated using the Mann-Whitney test comparing bezafibrate-treated OAC groups to the DMSO-treated OAC control group, and values are expressed as means + SEM [A-B]. Each point represents a patient [A, B, and D]. *Abbreviations: BMI, Body Mass Index; KL score, Kellgren-Lawrence score; RBC, Red Blood Cells; WBC, White Blood Cells*.

Given this dual response, we next investigated clinical data to identify potential discriminant parameters associated with bezafibrate treatment responsiveness. Clinical parameters included demographic (sex, age, BMI), serological (e.g., platelets, monocytes), and radiological (Kellgren-Lawrence score and surgical sampling side) features. The radar plot presented in **Figure 3C** provides an overview of these clinical parameters, showing a similar profile between groups for sex and age, whereas BMI, red blood cell counts and neutrophils levels appeared more divergent. Although univariate comparisons of these parameters between responders and non-responders groups showed no significant differences (**Supplementary Table 2**), multivariate Partial Least Squares-Discriminant Analysis (PLS-DA) revealed a clear separation between both groups (**Figure 3D**), with lower red and white blood cell counts, and lower BMI contributing most to the discrimination of responders as illustrated by the Variable Importance in Projection (VIP) scores (**Figure 3E**). Such multivariate PLS-DA analysis could provide a basis for predicting OAC responsiveness to bezafibrate.

## DISCUSSION

Mitochondrial dysfunction in OA chondrocytes has recently gained attention and is now recognized as a feature of the disease that can contribute to its progression [9–11]. Despite evidence of mitochondrial dysfunction and the potential benefits of restoring it, particularly through the promotion of OxPhos, yet no mitochondria-based therapies have yet been developed to treat OA. Indeed, current approaches such as mitochondrial transplantation have shown promising efficacy but still face challenges regarding their long-term safety and clinical efficacy [26–28]. Moreover, mitochondrial dysfunction in OA chondrocytes remains insufficiently characterized. Here, we revealed that OAC exhibit a higher number of mitochondria along with OxPhos dysfunction, and that this dysfunction can be partially restored by bezafibrate in a subgroup of patients.

FTIR, a non-destructive technique that can detect overall chemical variations, was first used to explore tissue-level changes in damaged compared to undamaged OA cartilage zones. This approach has already been used in the context of OA cartilage, to evaluate changes in collagen and proteoglycan content in extracellular matrix composition [36,37]. Interestingly, FTIR can also provide information on chondrocyte glucose metabolism by assessing glycogen and lactic acid content [32]. By assessing the full cartilage thickness, highly heterogeneous proteoglycan, glycogen and lactic acid profiles were observed between patients. Nevertheless, proteoglycan and glycogen deposits levels negatively correlated with OARSI scores. FTIR partially corroborated with STEM results at the cellular level, which revealed a trend toward decreased glycogen deposits in OA chondrocytes extracted from the damaged cartilage, when compared to undamaged areas. Further investigations will be required to define more precise regions of interest and distinguish signals from chondrocytes or their pericellular zone and their interterritorial ECM, as well as to differentiate cartilage layers, which could help uncover differences between experimental groups.

In line with what was observed in chondrocytes from damaged versus undamaged OA cartilage, OAC imaged by STEM displayed a significant decline in glycogen compared to NC. Knowing that glucose is the main energy source for chondrocytes, OAC may mobilize it from glycogen storage through glycogenolysis. This mechanism could serve as an adaptive response to meet energy demands, as previously observed during chondrogenesis [38]. OAC may indeed require this glucose to counterbalance the reduction in energy production efficiency due to the metabolic shift toward glycolysis at the expense of OxPhos, which is normally more efficient for energy production. Since this shift could be driven by initial mtDNA mutations leading to impaired

OxPhos in OA chondrocytes [39,40], we also examined potential changes in mitochondria by STEM. While mitochondria size was similar between NC and OAC, it tended to decrease in chondrocytes from more damaged compared to undamaged OA cartilage zones, which could be related to higher levels of fission processes, as already described in human OA chondrocytes in vitro [12]. However, in our settings, mitochondria shape was not modified between experimental conditions. Further analyses using deep learning-based segmentation of mitochondria on STEM images, and ultrastructural characterization approaches would enable a more precise assessment of mitochondrial morphological parameters [41–43].

Mitochondria number was not significantly affected in chondrocytes from damaged as compared to undamaged OA cartilage, whereas in OAC compared to NC controls, mitochondria number, as well as mtDNA copy number were significantly increased. Interestingly, a similar increase in mtDNA copy number has also been reported in primary human OA chondrocytes, thus further supporting our findings [44]. However, it should be noted that the number of mtDNA copies does not equate directly to the number of mitochondria, but rather provides an indirect estimation of mitochondrial content, for which STEM could offer a more accurate representation. However, using STEM, different studies reported opposite results with increased glycogen levels and decreased mitochondria number in primary human OA chondrocytes [23,45,46]. This discrepancy could be explained by different anatomical locations, as well as by isolation and culture steps, as passages could affect the number of mitochondria in primary chondrocytes [47]. In this study, we cannot completely exclude the possibility that the observed changes are attributable to differences in the anatomical locations and ages of the donors. Nevertheless, we have previously used these cells to study OA-related pathological changes and demonstrated their reliability compared with other cell types using public datasets [25].

Overall, we believe that the results presented herein, obtained using a multimodal approach combining FTIR, STEM, and mtDNA copy number quantification in native cartilage from two different pathological models, more accurately reflect the reality of glycogen contents and mitochondrial number in OA chondrocytes as compared to what has been previously reported in isolated and cultured cells in vitro. Both glycogen deposits reduction and mitochondrial number increase could be hypothesized as a compensatory response to balance bioenergetics failure and initial mitochondrial dysfunction.

Functional assays further revealed that OAC exhibit mitochondrial respiratory failure, including lower values of basal respiration, maximal respiration, and ATP production, as well as a reduced contribution of OxPhos for ATP production, compared to NC. This was consistent with existing literature reporting similar results in OA chondrocyte models assessed using the same real-time cellular metabolic assay approach [17,25,48,49] or with metabolomics and C13 fluxomics analyses [23]. In this context, we evaluated the potential of bezafibrate, an approved PPAR agonist drug used to treat dyslipidemia, which has also been shown to have promising effects in promoting OxPhos activity in other contexts, both in vitro, in vivo, and in clinical studies [34,50,51]. Patient-derived OAC could be classified as responders or non-responders to the 1 µM bezafibrate dose for 24 h. Such OxPhos-enhancing effects of bezafibrate could occur through various mechanisms. As a pan-PPAR agonist, bezafibrate promotes mitochondrial biogenesis by activating PGC-1α [50,52]. This increases the mitochondrial pool and could support overall cellular OxPhos. Another possible mechanism could involve enhanced fatty acid oxidation, which may lead to increased levels of acetyl-CoA that fuel the TCA cycle and, consequently, OxPhos [53].

While further investigation is needed to elucidate the underlying mechanism of action, bezafibrate OAC responsiveness and clinical data were integrated using a PLS-DA multivariate analysis, which helped separate the responders and non-responders groups. Although this analysis was supervised and limited to a small sample size, it provided initial insights that could contribute to the development of a future predictive model to identify a typical patient profile responsive to mitochondria-based therapies such as bezafibrate.

Altogether, our study benefits from a multimodal approach and reveals that OAC display altered glucose metabolism illustrated by a decreased glycogen deposits surface, increased mitochondrial number, and OxPhos dysfunction. This study also highlighted the ability of bezafibrate to improve OxPhos parameters in a specific subgroup of patient-derived OAC. Further investigation is now needed to determine the potential mechanism by which bezafibrate promotes OxPhos in OAC. Although no direct correlation could be established between the metabolic features investigated by FTIR and STEM, and the responsiveness to bezafibrate assessed by real-time cellular metabolic assays, integrating these approaches in future studies may yield a more comprehensive understanding of patient heterogeneity and responses to therapeutic molecules. To better explore metabolism processes at the cellular level, a single-cell metabolomics approach, such as the SCENITH technology [54] could offer a promising avenue to decipher OAC metabolic diversity, identify relevant subpopulations, and evaluate their enrichment across distinct patient subgroups. Overall, our study emphasizes the importance of deciphering OA patient metabolic diversity to develop effective mitochondria-based therapies, such as bezafibrate, in the future.

## Author contributions

Conceptualization: LD, RG, MAB and CV, Methodology: LD, RG, AD, GM, JV, MAB and CV, Investigation: LD, RG, SH, NG, GM, JV, CV and MAB, Sample providers: AH and DW, Supervision: MAB and CV, Writing-original draft: LD, MAB and CV, Writing-review & editing: LD, RG, AD, JV, GM, JG, MAB and CV.

## Acknowledgments

The authors thank Claire Pecqueur (CRCI2NA, Nantes Université, INSERM U1307, CNRS 6075, Nantes, France) for her help with Seahorse technology, with financial support from ITMO Cancer of Aviesan within the framework of the 2021-2030 Cancer Control Strategy. The authors thank Nina Bon for her help in performing and analyzing mtDNA quantification experiments. The authors also thank Manon Bauerheim for her help in collecting clinical data. The authors acknowledge the SC3M facility and the Sc4Bio from the Inserm/Nantes Université/ONIRIS UMR1229 RMeS Laboratory, SFR Bonamy and SFR ICAT. The authors also acknowledge the HiMolA facility from the Inserm/Université d’Angers/Nantes Université/ONIRIS UMR1229 RMeS Laboratory. They also acknowledge the MicroPICell core facility (SFR Bonamy, BioCore, Inserm UMS 016, CNRS UAR 3556, Nantes, France), member of the Scientific Interest Group (GIS) Biogenouest, IBISA, and the national infrastructure France-Bioimaging supported by the French national research agency (ANR-10-INBS-04).

## REFERENCES

[1] Steinmetz JD, Culbreth GT, Haile LM, Rafferty Q, Lo J, Fukutaki KG, et al. Global, regional, and national burden of osteoarthritis, 1990–2020 and projections to 2050: a systematic analysis for the Global Burden of Disease Study 2021. The Lancet Rheumatology 2023;5:e508–22. 10.1016/S2665-9913(23)00163-7.

[2] Kolasinski SL, Neogi T, Hochberg MC, Oatis C, Guyatt G, Block J, et al. 2019 American College of Rheumatology/Arthritis Foundation Guideline for the Management of Osteoarthritis of the Hand, Hip, and Knee. Arthritis Care Res (Hoboken) 2020;72:149–62. 10.1002/acr.24131.

[3] Leifer VP, Katz JN, Losina E. The burden of OA-health services and economics. Osteoarthritis Cartilage 2022;30:10–6. 10.1016/j.joca.2021.05.007.

[4] Oo WM, Little C, Duong V, Hunter DJ. The Development of Disease-Modifying Therapies for Osteoarthritis (DMOADs): The Evidence to Date. Drug Des Devel Ther 2021;15:2921–45. 10.2147/DDDT.S295224.

[5] Dell’Isola A, Allan R, Smith SL, Marreiros SSP, Steultjens M. Identification of clinical phenotypes in knee osteoarthritis: a systematic review of the literature. BMC Musculoskeletal Disorders 2016;17:425. 10.1186/s12891-016-1286-2.

[6] Mobasheri A, van Spil WE, Budd E, Uzieliene I, Bernotiene E, Bay-Jensen A-C, et al. Molecular taxonomy of osteoarthritis for patient stratification, disease management and drug development: biochemical markers associated with emerging clinical phenotypes and molecular endotypes. Current Opinion in Rheumatology 2019;31:80. 10.1097/BOR.0000000000000567.

[7] Mobasheri A, Loeser R. Clinical phenotypes, molecular endotypes and theratypes in OA therapeutic development. Nat Rev Rheumatol 2024;20:525–6. 10.1038/s41584-024-01126-4.

[8] López-Otín C, Blasco MA, Partridge L, Serrano M, Kroemer G. Hallmarks of aging: An expanding universe. Cell 2023;186:243–78. 10.1016/j.cell.2022.11.001.

[9] Blanco FJ, Rego I, Ruiz-Romero C. The role of mitochondria in osteoarthritis. Nat Rev Rheumatol 2011;7:161–9. 10.1038/nrrheum.2010.213.

[10] Zheng L, Zhang Z, Sheng P, Mobasheri A. The role of metabolism in chondrocyte dysfunction and the progression of osteoarthritis. Ageing Res Rev 2021;66:101249. 10.1016/j.arr.2020.101249.

[11] Tan S, Sun Y, Li S, Wu H, Ding Y. The impact of mitochondrial dysfunction on osteoarthritis cartilage: current insights and emerging mitochondria-targeted therapies. Bone Res 2025;13:77. 10.1038/s41413-025-00460-x.

[12] Ansari MY, Novak K, Haqqi TM. ERK1/2-mediated activation of DRP1 regulates mitochondrial dynamics and apoptosis in chondrocytes. Osteoarthritis Cartilage 2022;30:315–28. 10.1016/j.joca.2021.11.003.

[13] Wang Y, Zhao X, Lotz M, Terkeltaub R, Liu-Bryan R. Mitochondrial biogenesis is impaired in osteoarthritis chondrocytes but reversible via peroxisome proliferator-activated receptor γ coactivator 1α. Arthritis Rheumatol 2015;67:2141–53. 10.1002/art.39182.

[14] Ansari MY, Khan NM, Ahmad I, Haqqi TM. Parkin clearance of dysfunctional mitochondria regulates ROS levels and increases survival of human chondrocytes. Osteoarthritis Cartilage 2018;26:1087–97. 10.1016/j.joca.2017.07.020.

[15] Ma Y, Pang Y, Cao R, Zheng Z, Zheng K, Tian Y, et al. Targeting Parkin-regulated metabolomic change in cartilage in the treatment of osteoarthritis. iScience 2024;27. 10.1016/j.isci.2024.110597.

[16] Shin HJ, Park H, Shin N, Kwon HH, Yin Y, Hwang J-A, et al. Pink1-Mediated Chondrocytic Mitophagy Contributes to Cartilage Degeneration in Osteoarthritis. J Clin Med 2019;8:1849. 10.3390/jcm8111849.

[17] Ohashi Y, Takahashi N, Terabe K, Tsuchiya S, Kojima T, Knudson CB, et al. Metabolic reprogramming in chondrocytes to promote mitochondrial respiration reduces downstream features of osteoarthritis. Sci Rep 2021;11:15131. 10.1038/s41598-021-94611-9.

[18] Maneiro E, Martín MA, de Andres MC, López-Armada MJ, Fernández-Sueiro JL, del Hoyo P, et al. Mitochondrial respiratory activity is altered in osteoarthritic human articular chondrocytes. Arthritis & Rheumatism 2003;48:700–8. 10.1002/art.10837.

[19] Walsh DA, McWilliams DF, Turley MJ, Dixon MR, Fransès RE, Mapp PI, et al. Angiogenesis and nerve growth factor at the osteochondral junction in rheumatoid arthritis and osteoarthritis. Rheumatology (Oxford) 2010;49:1852–61. 10.1093/rheumatology/keq188.

[20] Vander Heiden MG, Cantley LC, Thompson CB. Understanding the Warburg Effect: The Metabolic Requirements of Cell Proliferation. Science 2009;324:1029–33. 10.1126/science.1160809.

[21] Yang X, Chen W, Zhao X, Chen L, Li W, Ran J, et al. Pyruvate Kinase M2 Modulates the Glycolysis of Chondrocyte and Extracellular Matrix in Osteoarthritis. DNA Cell Biol 2018;37:271–7. 10.1089/dna.2017.4048.

[22] Li HM, Guo HL, Xu C, Liu L, Hu SY, Hu ZH, et al. Inhibition of glycolysis by targeting lactate dehydrogenase A facilitates hyaluronan synthase 2 synthesis in synovial fibroblasts of temporomandibular joint osteoarthritis. Bone 2020;141:115584. 10.1016/j.bone.2020.115584.

[23] Wu X, Liyanage C, Plan M, Stark T, McCubbin T, Barrero RA, et al. Dysregulated energy metabolism impairs chondrocyte function in osteoarthritis. Osteoarthritis and Cartilage 2023;31:613–26. 10.1016/j.joca.2022.11.004.

[24] Cillero-Pastor B, Rego-Pérez I, Oreiro N, Fernandez-Lopez C, Blanco FJ. Mitochondrial respiratory chain dysfunction modulates metalloproteases -1, -3 and -13 in human normal chondrocytes in culture. BMC Musculoskeletal Disorders 2013;14:235. 10.1186/1471-2474-14-235.

[25] Defois A, Bon N, Charpentier A, Georget M, Gaigeard N, Blanchard F, et al. Osteoarthritic chondrocytes undergo a glycolysis-related metabolic switch upon exposure to IL-1b or TNF. Cell Commun Signal 2023;21:137. 10.1186/s12964-023-01150-z.

[26] Yu M, Wang D, Chen X, Zhong D, Luo J. BMSCs-derived Mitochondria Improve Osteoarthritis by Ameliorating Mitochondrial Dysfunction and Promoting Mitochondrial Biogenesis in Chondrocytes. Stem Cell Rev and Rep 2022;18:3092–111. 10.1007/s12015-022-10436-7.

[27] Lee AR, Woo JS, Lee S-Y, Na HS, Cho K-H, Lee YS, et al. Mitochondrial Transplantation Ameliorates the Development and Progression of Osteoarthritis. Immune Netw 2022;22:e14. 10.4110/in.2022.22.e14.

[28] Vega-Letter AM, García-Guerrero C, Yantén-Fuentes L, Pradenas C, Herrera-Luna Y, Lara- Barba E, et al. Safety and efficacy of mesenchymal stromal cells mitochondria transplantation as a cell-free therapy for osteoarthritis. J Transl Med 2025;23:26. 10.1186/s12967-024-05945-7.

[29] Chen P, Liu X, Gu C, Zhong P, Song N, Li M, et al. A plant-derived natural photosynthetic system for improving cell anabolism. Nature 2022;612:546–54. 10.1038/s41586-022-05499-y.

[30] Pritzker KPH, Gay S, Jimenez SA, Ostergaard K, Pelletier J-P, Revell PA, et al. Osteoarthritis cartilage histopathology: grading and staging. Osteoarthritis Cartilage 2006;14:13–29. 10.1016/j.joca.2005.07.014.

[31] Rieppo L, Närhi T, Helminen HJ, Jurvelin JS, Saarakkala S, Rieppo J. Infrared spectroscopic analysis of human and bovine articular cartilage proteoglycans using carbohydrate peak or its second derivative. J Biomed Opt 2013;18:097006. 10.1117/1.JBO.18.9.097006.

[32] Hackett MJ, Sylvain NJ, Hou H, Caine S, Alaverdashvili M, Pushie MJ, et al. Concurrent Glycogen and Lactate Imaging with FTIR Spectroscopy To Spatially Localize Metabolic Parameters of the Glial Response Following Brain Ischemia. Anal Chem 2016;88:10949–56. 10.1021/acs.analchem.6b02588.

[33] Jeroundi N, Roy C, Basset L, Pignon P, Preisser L, Blanchard S, et al. Glycogenesis and glyconeogenesis from glutamine, lactate and glycerol support human macrophage functions. EMBO Rep 2024;25:5383–407. 10.1038/s44319-024-00278-4.

[34] Douiev L, Sheffer R, Horvath G, Saada A. Bezafibrate Improves Mitochondrial Fission and Function in DNM1L-Deficient Patient Cells. Cells 2020;9:301. 10.3390/cells9020301.

[35] Augustyniak J, Lenart J, Gaj P, Kolanowska M, Jazdzewski K, Stepien PP, et al. Bezafibrate Upregulates Mitochondrial Biogenesis and Influence Neural Differentiation of Human-Induced Pluripotent Stem Cells. Mol Neurobiol 2019;56:4346–63. 10.1007/s12035-018-1368-2.

[36] Oinas J, Rieppo L, Finnilä M a. J, Valkealahti M, Lehenkari P, Saarakkala S. Imaging of Osteoarthritic Human Articular Cartilage using Fourier Transform Infrared Microspectroscopy Combined with Multivariate and Univariate Analysis. Sci Rep 2016;6:30008. 10.1038/srep30008.

[37] Linus A, Ebrahimi M, Turunen MJ, Saarakkala S, Joukainen A, Kröger H, et al. High-resolution infrared microspectroscopic characterization of cartilage cell microenvironment. Acta Biomaterialia 2021;134:252–60. 10.1016/j.actbio.2021.08.001.

[38] Schajowicz F, Cabrini RL. Histochemical Studies on Glycogen in Normal Ossification and Calcification. The Journal of Bone & Joint Surgery 1958;40:1081–92.

[39] Chang M-C, Hung S-C, Chen WY-K, Chen T-L, Lee C-F, Lee H-C, et al. Accumulation of mitochondrial DNA with 4977-bp deletion in knee cartilage--an association with idiopathic osteoarthritis. Osteoarthritis Cartilage 2005;13:1004–11. 10.1016/j.joca.2005.06.011.

[40] Kim J, Xu M, Xo R, Mates A, Wilson GL, Pearsall AW, et al. Mitochondrial DNA damage is involved in apoptosis caused by pro-inflammatory cytokines in human OA chondrocytes. Osteoarthritis and Cartilage 2010;18:424–32. 10.1016/j.joca.2009.09.008.

[41] Guay MD, Emam ZAS, Anderson AB, Aronova MA, Pokrovskaya ID, Storrie B, et al. Dense cellular segmentation for EM using 2D-3D neural network ensembles. Sci Rep 2021;11:2561. 10.1038/s41598-021-81590-0.

[42] Conrad R, Narayan K. Instance segmentation of mitochondria in electron microscopy images with a generalist deep learning model trained on a diverse dataset. Cell Syst 2023;14:58–71.e5. 10.1016/j.cels.2022.12.006.

[43] Jang C, Lee H, Yoo J, Yoon H. Deep learning-driven automated mitochondrial segmentation for analysis of complex transmission electron microscopy images. Sci Rep 2025;15:19076. 10.1038/s41598-025-03311-1.

[44] Dalmao-Fernández A, Hermida-Gómez T, Lund J, Vazquez-Mosquera ME, Rego-Pérez I, Garesse R, et al. Mitochondrial DNA from osteoarthritic patients drives functional impairment of mitochondrial activity: a study on transmitochondrial cybrids. Cytotherapy 2021;23:399–410. 10.1016/j.jcyt.2020.08.010.

[45] Fioravanti A, Nerucci F, Annefeld M, Collodel G, Marcolongo R. Morphological and cytoskeletal aspects of cultivated normal and osteoarthritic human articular chondrocytes after cyclical pressure: a pilot study. Clin Exp Rheumatol 2003;21:739–46.

[46] Pascarelli NA, Collodel G, Moretti E, Cheleschi S, Fioravanti A. Changes in Ultrastructure and Cytoskeletal Aspects of Human Normal and Osteoarthritic Chondrocytes Exposed to Interleukin-1β and Cyclical Hydrostatic Pressure. Int J Mol Sci 2015;16:26019–34. 10.3390/ijms161125936.

[47] Chen Y, Yu Y, Wen Y, Chen J, Lin J, Sheng Z, et al. A high-resolution route map reveals distinct stages of chondrocyte dedifferentiation for cartilage regeneration. Bone Res 2022;10:38. 10.1038/s41413-022-00209-w.

[48] Arra M, Swarnkar G, Ke K, Otero JE, Ying J, Duan X, et al. LDHA-mediated ROS generation in chondrocytes is a potential therapeutic target for osteoarthritis. Nat Commun 2020;11:3427. 10.1038/s41467-020-17242-0.

[49] Terabe K, Ohashi Y, Tsuchiya S, Ishizuka S, Knudson CB, Knudson W. Chondroprotective effects of 4-methylumbelliferone and hyaluronan synthase-2 overexpression involve changes in chondrocyte energy metabolism. J Biol Chem 2019;294:17799–817. 10.1074/jbc.RA119.009556.

[50] Komen JC, Thorburn DR. Turn up the power – pharmacological activation of mitochondrial biogenesis in mouse models. British Journal of Pharmacology 2014;171:1818–36. 10.1111/bph.12413.

[51] Steele H, Gomez-Duran A, Pyle A, Hopton S, Newman J, Stefanetti RJ, et al. Metabolic effects of bezafibrate in mitochondrial disease. EMBO Mol Med 2020;12:e11589. 10.15252/emmm.201911589.

[52] Grings M, Moura AP, Parmeggiani B, Pletsch JT, Cardoso GMF, August PM, et al. Bezafibrate prevents mitochondrial dysfunction, antioxidant system disturbance, glial reactivity and neuronal damage induced by sulfite administration in striatum of rats: Implications for a possible therapeutic strategy for sulfite oxidase deficiency. Biochimica et Biophysica Acta (BBA) - Molecular Basis of Disease 2017;1863:2135–48. 10.1016/j.bbadis.2017.05.019.

[53] Djouadi F, Aubey F, Schlemmer D, Ruiter JPN, Wanders RJA, Strauss AW, et al. Bezafibrate increases very-long-chain acyl-CoA dehydrogenase protein and mRNA expression in deficient fibroblasts and is a potential therapy for fatty acid oxidation disorders. Hum Mol Genet 2005;14:2695–703. 10.1093/hmg/ddi303.

[54] Argüello RJ, Combes AJ, Char R, Gigan J-P, Baaziz AI, Bousiquot E, et al. SCENITH: A Flow Cytometry-Based Method to Functionally Profile Energy Metabolism with Single-Cell Resolution. Cell Metab 2020;32:1063–1075.e7. 10.1016/j.cmet.2020.11.007.

