## Supplementary_information for "Multimodal analysis of osteoarthritic chondrocytes reveals mitochondrial alterations and patient-specific OxPhos response to bezafibrate"

**Supplementary Table 1. Demographic parameters of OA patients according to the different techniques used.**

| | **FTIR study** (N=6) | **STEM study** (N=5) | **Real-time cellular metabolic assay (Seahorse) study** | | | **p-values** ***FTIR study patients vs STEM study patients vs Seahorse study patients*** (Chi² or Kruskal-Wallis, as appropriate**)** |
|---|---|---|---|---|---|---|
| | | | All (N=14) | Responders (N=7) | Non-responders (N=7) | |
| **Female,** % (N) | **33%** (2) | **60%** (3) | **71.43%** (10) | **71.43%** (5) | **71.43%** (5) | 0.5671 ns |
| **Age,** (years)**,** mean (SD) | **67.17** (6.15) | **70.80** (10.28) | **71.28** (8.61) | **71.42** (7.93) | **71.14** (9.88) | 0.4693 ns |
| **BMI,** mean (SD) | **32.50** (6.30) | **31.64** (9.78) | **31.49** (6.21) | **30.60** (4.59) | **32.39** (7.77) | 0.9861 ns |
| **KL score,** mean (SD) | **3.67** (0.52) | **3.80** (0.45) | **3.43** (0.94) | **3.71** (0.49) | **3.14** (1.21) | 0.7490 ns |

*Abbreviations: BMI, Body Mass Index; FTIR, Fourier-Transform Infrared Spectroscopy; KL, Kellgren-Lawrence; STEM, Scanning Transmission Electron Microscopy.*

**Supplementary Table 2. Demographic parameters of OA patients according to the OAC responsiveness to bezafibrate in real-time cellular metabolic assays (Seahorse).**

| | **Non-responders (N=7)** | **Responders (N=7)** | **p-values** ***Responders vs Non-responders*** (Fisher's test or Mann-Whitney as appropriate) |
|---|---|---|---|
| **Female,** % (N) | **71.43%** (5) | **71.43%** (5) | >0,9999 ns |
| **Age,** (years), mean (SD) | **71.14** (9.89) | **71.42** (7.93) | 0,9732 ns |
| **BMI,** mean (SD) | **32.39** (7.78) | **30.60** (4.60) | 0.7873 ns |
| **KL score,** mean (SD) | **3.14** (1.36) | **3.71** (0.49) | 0.4860 ns |
| **Right sampling side,** % (N) | **71.43%** (5) | **71.43%** (5) | >0,9999 ns |
| **Hemoglobin,** (g/dL), mean (SD) | **12.68** (1.36) | **12.13** (1.16) | 0.5350 ns |
| **Hematocrit,** (%), mean (SD) | **38.30** (3.95) | **36.14** (4.38) | 0.1981 ns |
| **Red blood cells,** (T/L), mean (SD) | **4.42** (0.67) | **3.91** (0.36) | 0.1055 ns |
| **White blood cells,** (g/L), mean (SD) | **10.29** (7.01) | **11.30** (4.75) | 0.5350 ns |
| **Platelets,** (g/L), mean (SD) | **210.29** (38.10) | **219.57** (26.93) | >0,9999 ns |
| **Neutrophils,** (g/L), mean (SD) | **6.06** (3.58) | **8.49** (3.52) | 0.1212 ns |
| **Lymphocytes,** (g/L), mean (SD) | **3.15** (3.92) | **2.02** (2.30) | 0.1544 ns |
| **Monocytes,** (g/L), mean (SD) | **0.68** (0.24) | **1.56** (1.79) | 0.2570 ns |
| **Eosinophils,** (g/L), mean (SD) | **0.06** (0.07) | **0.04** (0.06) | 0.4231 ns |
| **Basophils,** (g/L), mean (SD) | **0.03** (0.02) | **0.02** (0.02) | 0.3881 ns |

*Abbreviations: BMI, Body Mass Index; KL: Kellgren-Lawrence.*

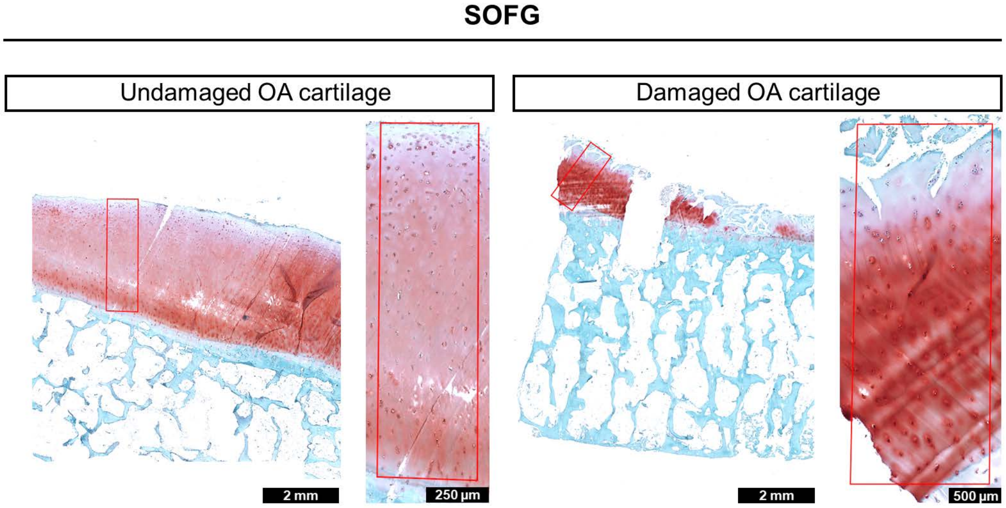


**Supplementary Figure 1.** Representative Safranin O/Fast Green staining of undamaged and damaged OA cartilage zones in the same patient (N=6). The representative images shown correspond to OA patient 4.

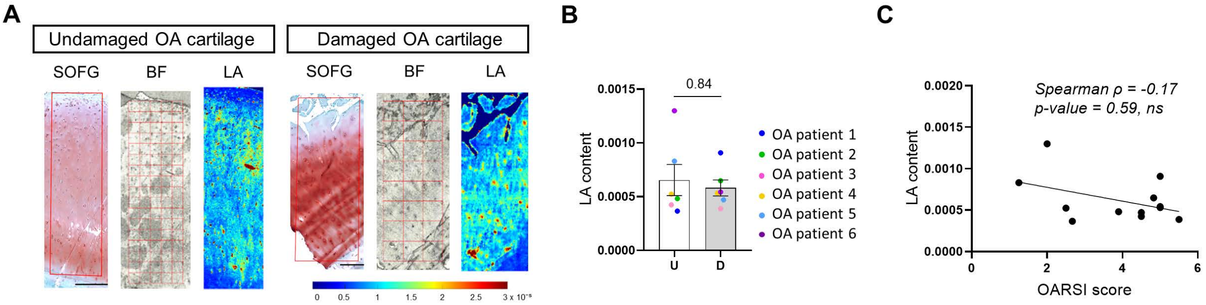


**Supplementary Figure 2. FTIR analyses of lactic acid content in undamaged (U) and damaged (D) OA cartilage. A.** FTIR spectroscopic imaging of lactic acid (LA). Safranin O/Fast Green (SOFG) staining was used to guide the selection of regions of interest (ROI) in the corresponding bright-field (BF) images for FTIR analysis. The representative images shown correspond to OA patient 4 (N=6). Scale bar, 250 µm for undamaged and 500 µm for damaged OA cartilage**. B.** Quantification of lactic acid content normalized to the amide I band measured by FTIR spectroscopy (N=6). Data are represented as the mean experimental spatial values from each patient's D or U zones. Values are expressed as means ± SEM. **C.** Spearman correlation between lactic acid content measured by FTIR in OA cartilage and OARSI scores. Mann-Whitney test comparing U and D was used for panel B and Spearman's correlation test was used for panel C. ns, non-significant. The corresponding p-values and Spearman's correlation coefficient (ρ, rho) are indicated on the appropriate graphs.

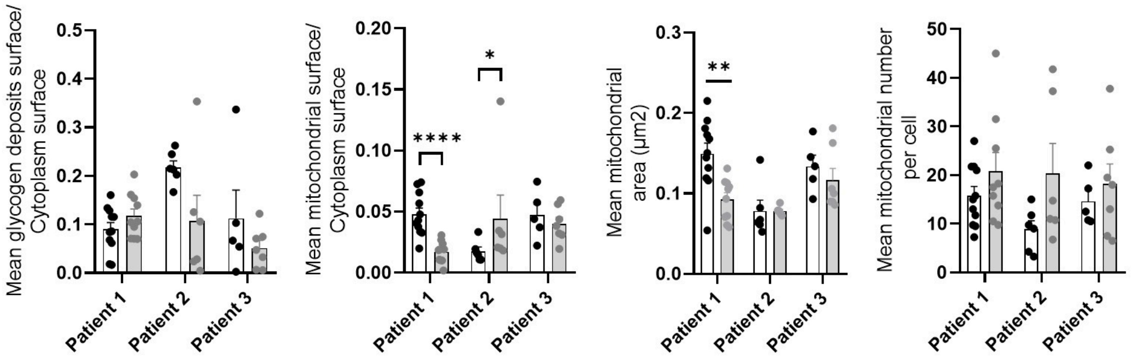


**Supplementary Figure 3.** Quantification of glycogen deposits and mitochondrial parameters from STEM images in chondrocytes from undamaged (U) compared to damaged (D) OA cartilage (N=3, n ≥ 6 cells). Data is presented for each patient. Each dot represents a cell. *p < 0.05, **p < 0.01, ***p < 0.001, ****p < 0.0001, Mann-Whitney test comparing undamaged and damaged OA cartilage. Values are expressed as means + SEM.

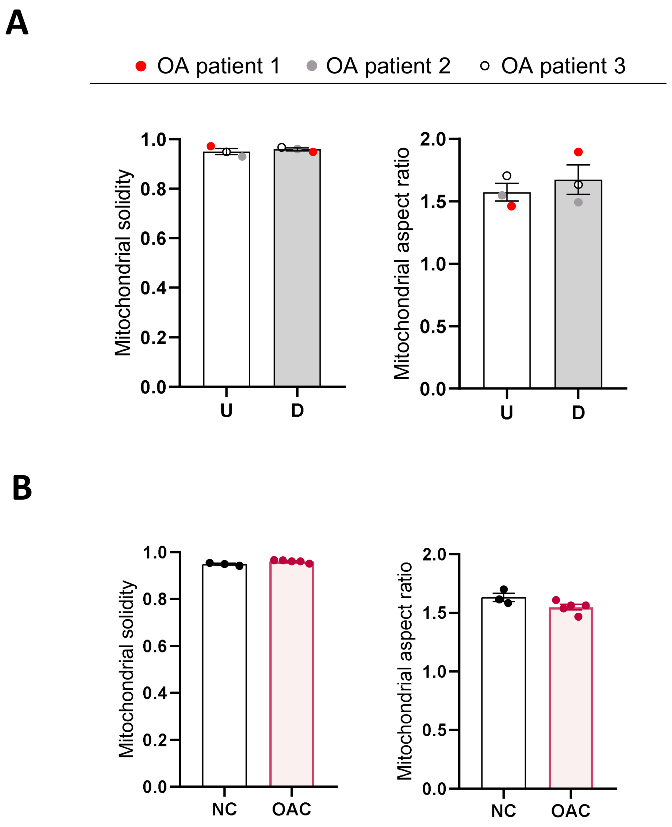


**Supplementary Figure 4.** Quantification of mitochondrial solidity and aspect ratio from STEM images. The panel A shows the data for chondrocytes from undamaged (U) versus damaged (D) OA cartilage (N=3, n ≥ 6 cells), while the panel B shows the data for NC (N=3, n ≥ 6 cells) compared to OAC (N=5, n ≥ 6 cells). Each dot represents a patient. Values are expressed as means ± SEM.

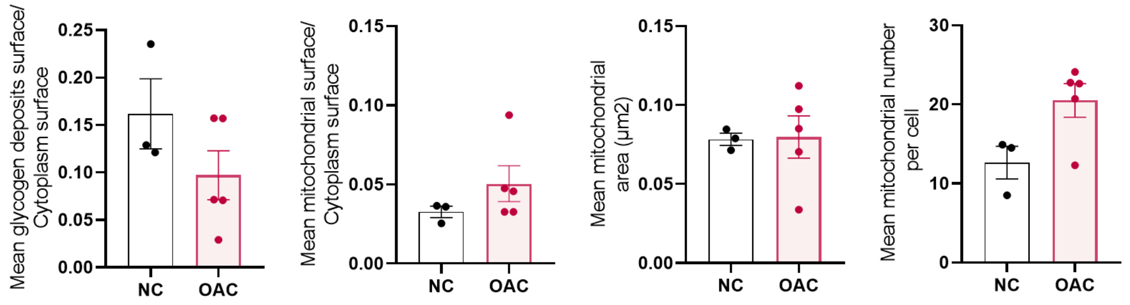


**Supplementary Figure 5.** Quantification of glycogen deposits and mitochondrial parameters from STEM images in OAC (N=5, n ≥ 6 cells) compared to NC (N=3, n ≥ 6 cells). Each dot represents a patient. Values are expressed as means ± SEM.

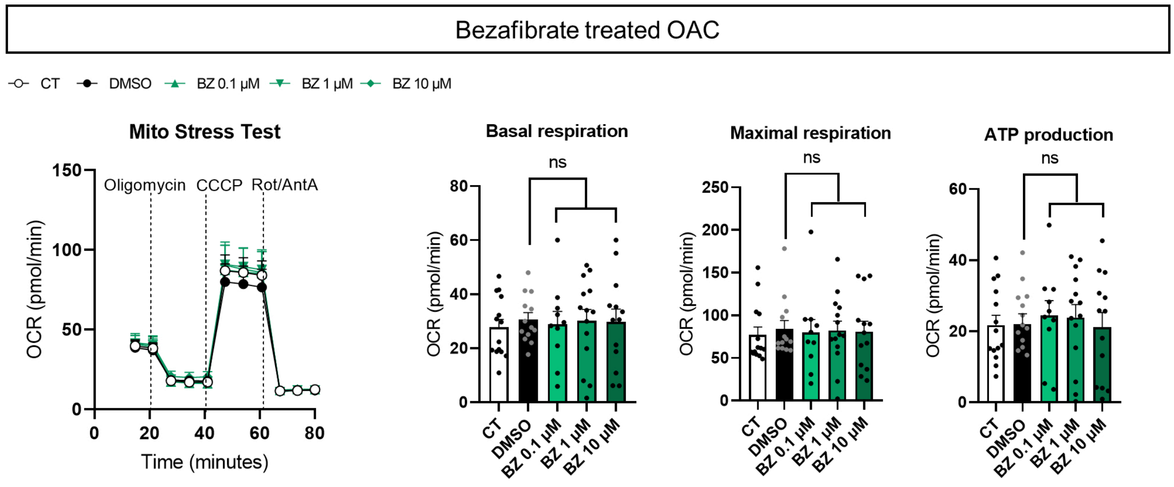


**Supplementary Figure 6.** Functional analysis of mitochondrial respiration in OAC after 24 h treatment with bezafibrate (0.1, 1 or 10 µM). OxPhos function was assessed using a Mito Stress Test (Seahorse). Oxygen consumption rate (OCR) values were used to quantify basal respiration, maximal respiration, and ATP production (N=14). Each dot represents a patient. Mann-Whitney test comparing DMSO and bezafibrate-treated groups. ns, non-significant. Values are expressed as means + SEM.
